# Co-expression analysis to identify key modules and hub genes associated with COVID19 in Platelets

**DOI:** 10.1101/2021.09.01.458644

**Authors:** Ahmed B. Alarabi, Attayeb Mohsen, Kenji Mizuguchi, Fatima Z. Alshbool, Fadi T. Khasawneh

**Author notes:** Correspondence: Fadi Khasawneh: Department of Pharmaceutical Sciences, Irma Lerma Rangel College of Pharmacy, Texas A&M University, Kingsville, Texas, USA,.

## Abstract

The severe acute respiratory syndrome corona virus 2 (SARS-CoV-2) is a highly contagious virus that causes a severe respiratory disease known as Corona virus disease 2019 (COVID19). Indeed, COVID19 increases the risk of cardiovascular occlusive/thrombotic events and is linked to poor outcomes. The pathophysiological processes underlying COVID19-induced thrombosis are complex, and remain poorly understood. To this end, platelets play important roles in regulating our cardiovascular system, including via contributions to coagulation and inflammation. There is an ample of evidence that circulating platelets are activated in COVID19 patients, which is a primary driver of the thrombotic outcome observed in these patients. However, the comprehensive molecular basis of platelet activation in COVID19 disease remains elusive, which warrants more investigation. Hence, we employed gene co-expression network analysis combined with pathways enrichment analysis to further investigate the aforementioned issues. Our study revealed three important gene clusters/modules that were closely related to COVID19. Furthermore, enrichment analysis showed that these three modules were mostly related to platelet metabolism, protein translation, mitochondrial activity, and oxidative phosphorylation, as well as regulation of megakaryocyte differentiation, and apoptosis, suggesting a hyperactivation status of platelets in COVID19. We identified the three hub genes from each of three key modules according to their intramodular connectivity value ranking, namely: COPE, CDC37, CAPNS1, AURKAIP1, LAMTOR2, GABARAP MT-ND1, MT-ND5, and MTRNR2L12. Collectively, our results offer a new and interesting insight into platelet involvement in COVID19 disease at the molecular level, which might aid in defining new targets for treatment of COVID19–induced thrombosis.

**key points:**

- Co-expression analysis of platelet RNAseq from COVID19 patients show distinct clusters of genes (modules) that are highly correlated to COVID19 disease.
- Identifying these modules might help in understanding the mechanism of thrombosis in COVID19 patients

## Introduction

The coronavirus SARS-CoV-2 is a highly contagious infection that causes a severe respiratory disease known as COVID19. This disease that has reached a pandemic level, is impacting tens of millions of people worldwide. In the United States, there are around 34 million reported cases, over two million hospital admissions, and half a million deaths as of July 2021^1^. It is now known that COVID19-induced thrombosis increases the incidence of cardiovascular occlusive events in infected patients, a fact that has been reported in several studies^2–4^, Indeed, abnormal hemostasis responses were observed in COVID19 hospitalized patients, which was linked to poor prognosis^2,5,6^ In addition, studies have shown that COVID19 leads to increase in platelet activation through alterations of platelet transcriptome and proteome^7,8^. In this connection, it is now well established that platelets play roles beyond vascular hemostasis, including innate immunity and tumor metastasis^9^. Moreover, platelets were shown to be activated in the septic state, and antiplatelet therapy has been used as a strategy to prevent organ damage in sepsis^10^ To this end, evidence have indicated that viral infections are associated with coagulation disorders, and thrombotic cardiovascular events^11,12^, which is consistent with the thrombotic phenotype seen in COVID19 patients/SARS-CoV-2 viral infection. While there has been some progress, our understanding of the pathways that govern platelet participation in COVID19–induced thrombosis remains limited, but clearly warrants investigation.

To obtain a comprehensive insight into the pathogenesis of specific disease states, several computational and research methods have been developed^13^. Some of these approaches were employed to examine the potential gene networks, which are very instrumental to guide understanding of diseases and their mechanistic pathways. Notably, co-expression analysis is one such approach, which clusters genes into coexpressed groups known as modules. These genes that belong to the same module are thought to share functional properties ^14^. This approach relies on using graph theory concepts that allow researchers to understand in a systematic way the relations between the genes of a module and the phenotype based on the module eigingene^14^. In fact, co-expression using weighted correlation network analysis (WGCNA) has been used for analyzing a number of biological processes, including cancer^15,16^ and cognitive and mental disorders^17,18^. In short, gene networks provide the utility to move beyond individual-gene comparisons and comprehensively identify biologically meaningful relationships between gene products and phenotypes.

Previous studies on the mechanisms of thrombosis in COVID19 disease have primarily concentrated on specific patho-physiological functions, with relatively fewer studies identifying comprehensive regulatory networks. Therefore, in the present study, WGCNA was used to determine gene networks associated with COVID19 disease in platelets. PRJNA634489 dataset-which contained a total of 15 samples from COVID19 patients and health controls ^7^ was used in the present study. Three modules with the highest level of significance in correlation with COVID19 disease were identified, and the three genes with the highest intramodular connectivity were selected as the hub genes in the respective modules for COVID19. Gene enrichment analysis was also conducted to determine enrichments in the key modules. The results of this study may provide novel information/insight into the underlying mechanisms of COVID19 disease and may assist in the identification of potential biomarkers for diagnosis and targets for treatment.

## Methods

### Data Preprocessing and Differentially Expressed Genes Screening

RNAseq data were downloaded from BioProject accession #PRJNA634489^7^. Data comprised of ten COVID19 patients in addition to age- and sex-matched five healthy controls. The Kallisto program was employed for pseudoalignment of reads and quantification to obtain the counts and the transcript per million (TPM)^19^. Log2CPM (log transformed counts per million) was used for the differential expression analysis by employing Voom normalization^20^ and Limma R package^21^ TPM normalized and filtered to exclude low variance transcripts (≤ 0.001)^22^ was used for the weighted gene co-expression network analysis.

### Weighted Gene Coexpression Network Analysis

The weighted co-expression network was produced using R package “WCGNA” ^14^. To weight highly correlated genes, the soft thresholding power (*β*) was set at 12, and the minimal module size was set at 30. To define clusters of genes in the dataset, the adjacency matrix was used to calculate the topological overlap matrix (TOM), which shows the degree of overlap in shared neighbors between pairs of genes in the network. The resulting gene network was visualized as a heatmap.

### Screening for key modules and hub genes

Correlation between module eigengenes and the COVID19 status was calculated to identify key modules that have significant correlation. The correlation values were displayed within a heatmap. The modules that correlated with COVID19 most significantly were considered as the key modules. Gene significance (GS) was defined as the correlation between gene expression and the COVID19 status. Module membership (MM) was defined as the correlation between gene expression and each module’s eigengene, and intramodular connectivity (K.in), which measures how connected a given gene with respect to the genes of a particular module, was also calculated using WGCNA. Subsequently, the correlation between GS and MM as well as GS and k.in were examined to verify module-COVID19 status associations. The correlation analyses in this study were performed using the Pearson correlation as described in the “WGCNA” package^14^. All module genes were ranked according to their intramodular connectivity, and only the top three genes were selected as hub genes.

### Functional enrichment analysis of key modules

The genes in each key modules were extracted from the network and enrichment analysis was performed to further explore the functions of the respective modules. Targetmine^23^ which is a web-based integrative data analysis platform for target prioritisation and broad-based biological knowledge discovery-was used to perform Gene Ontology (GO) and Reactome pathway enrichment analysis. In this analysis, a benjamini hochberg adjusted P-value of 0.05 was set as the significance threshold to identify the most significant functional pathways/GO terms. Only top results of enriched terms are reported.

### Statistical and visualization tools

We used R statistical programming language^24^ version 4.1.0, with the following packages: “WGCNA”^14^ for coexpression analysis; “circlize”^25^ for chord diagram building; “ggplot2” for visualization ^26^; “Igraph” for network analysis^27^ and “ggraph”^28^ for network visualization.

## Results

### Construction of Co-expression Network

The transcript per million (TPM) gene expression dataset were filtered based on variance, and 7119 genes in the 15 samples of ten COVID19 patients and five healthy controls were used to construct the co-expression network. The results of cluster analysis of the samples are demonstrated in (Fig. 1A). To construct the network, a soft-threshold of 12 was used to obtain the approximate scale-free topology (Supplementary Fig. 1). Genes across the 15 samples were hierarchically clustered based on topological overlap (Fig. 1C & 1D). We identified 16 modules in which genes are coexpressed, random colors were assigned to the modules to distinguish between them. The size (number of genes/module) of each module is presented in (Fig 1B). To demonstrate how these modules were relatively distinctive, we plotted the network heatmap of 400 randomly selected genes based on topological overlap matrix dissimilarity and their cluster dendrogram (Fig. 2A) indicating relative independence among these clusters.

**Figure 1:**
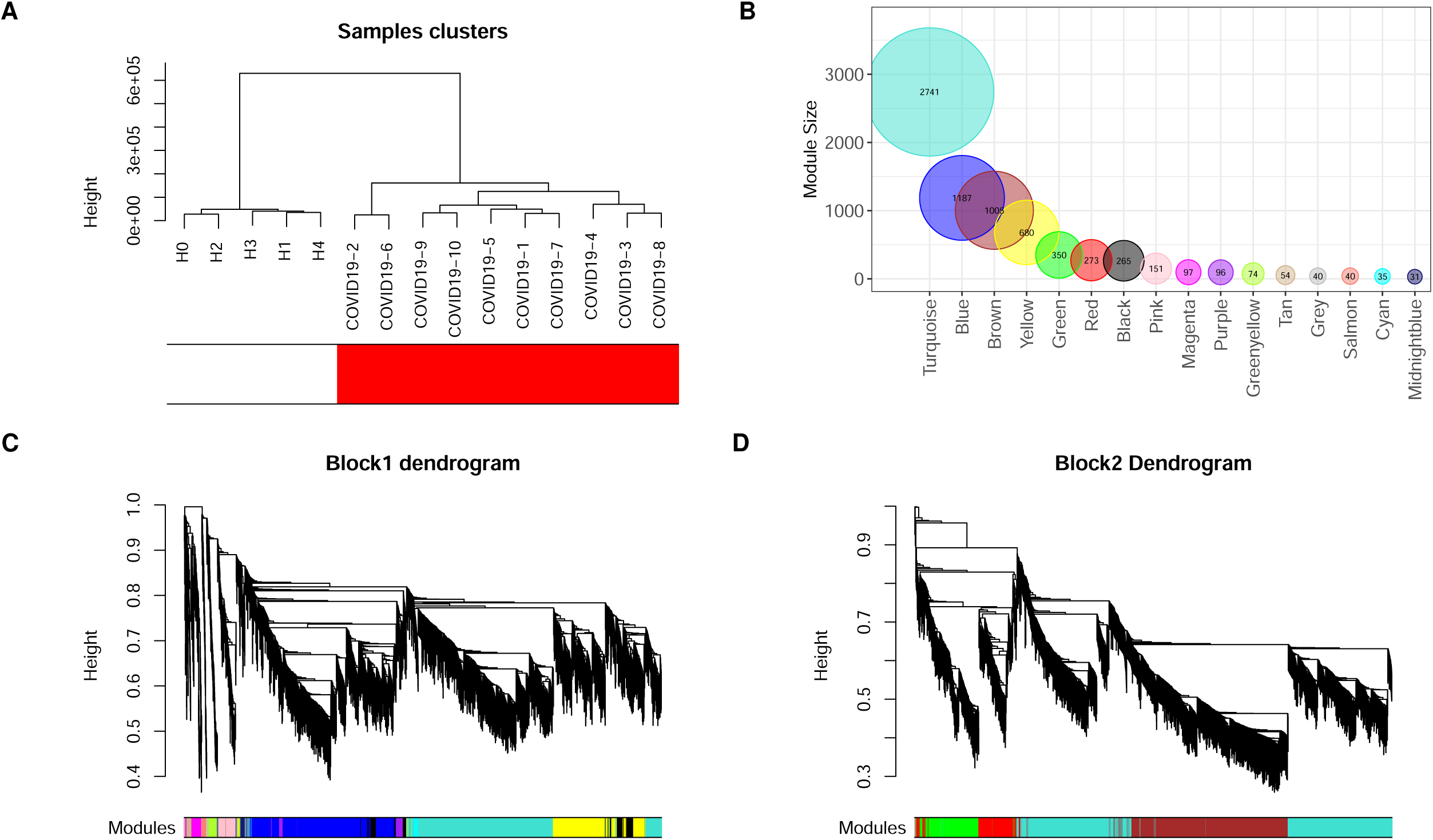
Construction of weighted co-expression network. (A) Sample dendrogram and trait heatmap. (B) Module size. (C and D) Cluster dendrogram block 1 and block 2. Each color represents one specific co-expression module, and branches above represent genes.

**Figure 2:**
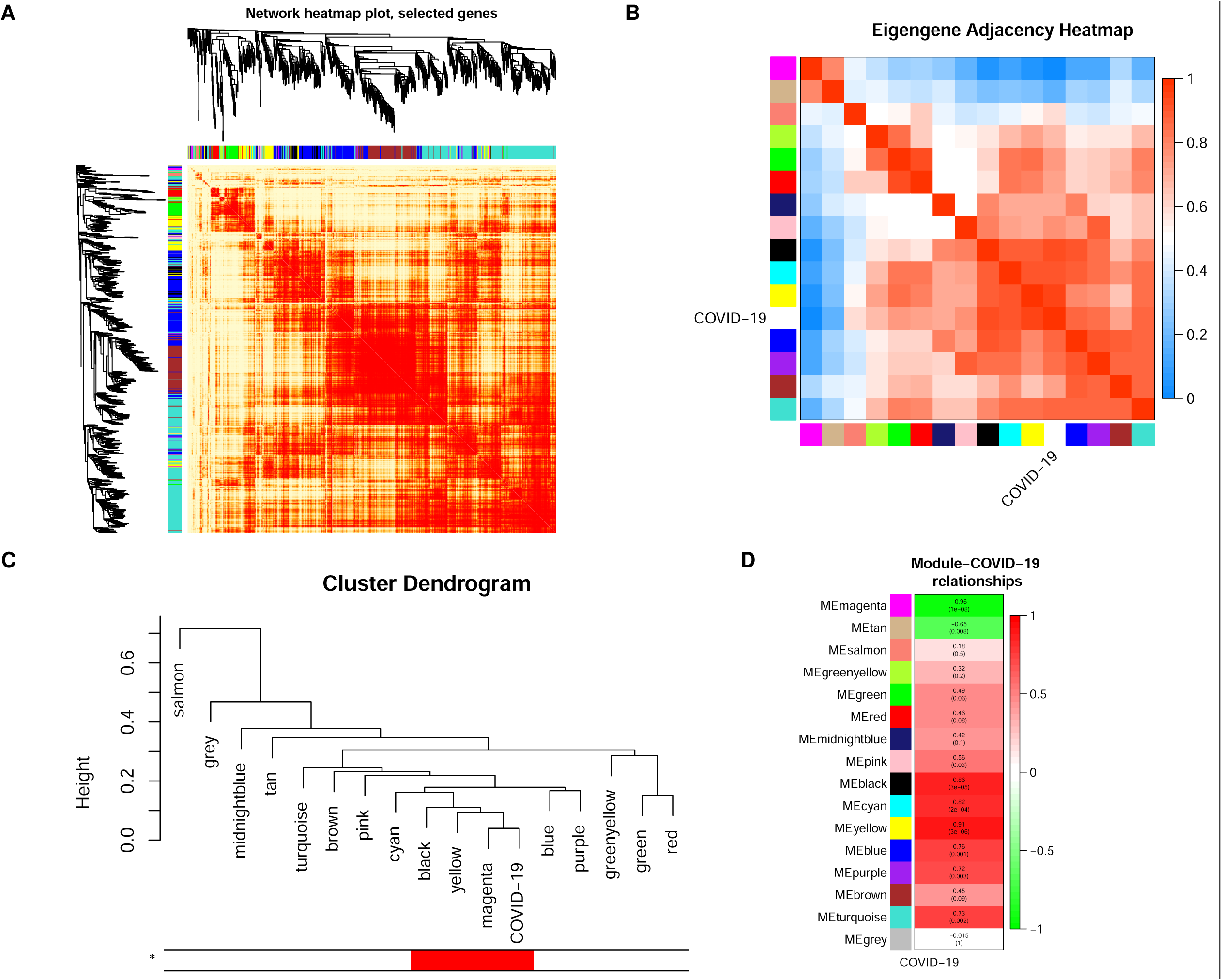
Co-expression module analysis. (A) Interaction of co-expression genes based on TOM dissimilarity and the cluster dendrogram of 1,000 randomly selected genes. The colors of the axes represent respective modules. The intensity of the yellow inside the heatmap represents the degree of overlap, with a darker yellow representing an increased overlap. (B) Eigengene adjacency heatmap of different gene co-expression modules. (C) An eigengene dendrogram identified groups of correlated modules. Heatmap of the correlation between COVID19 status and module eigengenes. Column corresponds to a clinical trait, and each row corresponds to a module. Each row contains the correlation coefficients which correspond to the cell color; green represents negative correlation and red represents positive correlation. The P-values are stated in the brackets.

### Correlation between modules and COVID19 disease status

To examine the relation of COVID19 status with the emerged modules, we built the eigengene adjacency matrix by calculating the correlation of the eigengenes matrix after inserting COVID19 status to the matrix. The heatmap (Fig. 2B) showed the modules’ relationship and the correlation between the modules namely black, cyan, yellow, blue, and magenta and COVID19 status.

### Identification of key modules in relationship to COVID19 disease status

To further determine the closest modules to COVID19 status, we re-clustered the eigengenes using single linkage method with absolute correlation as a distance function; the single linkage clustering algorithm looks for closest modules to form a cluster, then cluster them with the next nearest module progressively until one cluster is formed^29^. As demonstrated in Fig. 2C, the closest three modules to COVID19 status are magenta, yellow and black. Three essential measurements can help confirm the importance of the module to a specific trait, 1) Module membership (MM), which increases for a particular gene, when the module eigengene accurately represents this gene, 2) gene significance (GS) is measured by calculating the correlation of gene expression with the specific trait and 3) intramodular connectivity (K.in) for a gene within the module, reflecting the centrality of the gene to the module expression network. Based on WGCNA, if a gene is higher with GS, MM, and K.in, it is more meaningful to the clinical trait of interest ^30,31^. Explicitly, the higher the correlation between gene significance of genes in a module and their module membership, the higher its importance. Similarly, when the gene centrality in the network increases in parallel with gene significance, that also is strong evidence that key modules are essential in that trait. The correlations between gene significance and module membership as well as between gene significance and intramodular connectivity show that yellow, black, and magenta modules have the highest correlation values with a substantial difference to the next nearest module (Blue R=0.61) (Supplementary Fig. 2). For those reasons, we selected yellow, black, and magenta modules for further investigation and will refer to them using the term key modules.

### Key modules show high correlation to COVID19 disease status

The module-trait relationship was determined by correlating module eigengenes with COVID19 disease status to identify significant correlation. The yellow and the black modules exhibited the highest positive correlation (R=0.91; p-value=3 × 10^−6^, and R=0.86; p-value= 3 × 10^−5^, respectively; Fig. 2D). On the other hand, the magenta module (R=-0.96; p-value=1 × 10^−8^) exhibited the highest negative correlation (Fig. 2D). Therefore, these three modules were identified as key modules for COVID19 disease and its impact on platelets. The significant correlations between the different GS, MM, and K.in for COVID19 are illustrated in (Fig. 3A & 3B). We also showed the GS, MM, and K.in of the green module that showed the low correlation to COVID19 disease status (Fig. 3A & 3B).

**Figure 3:**
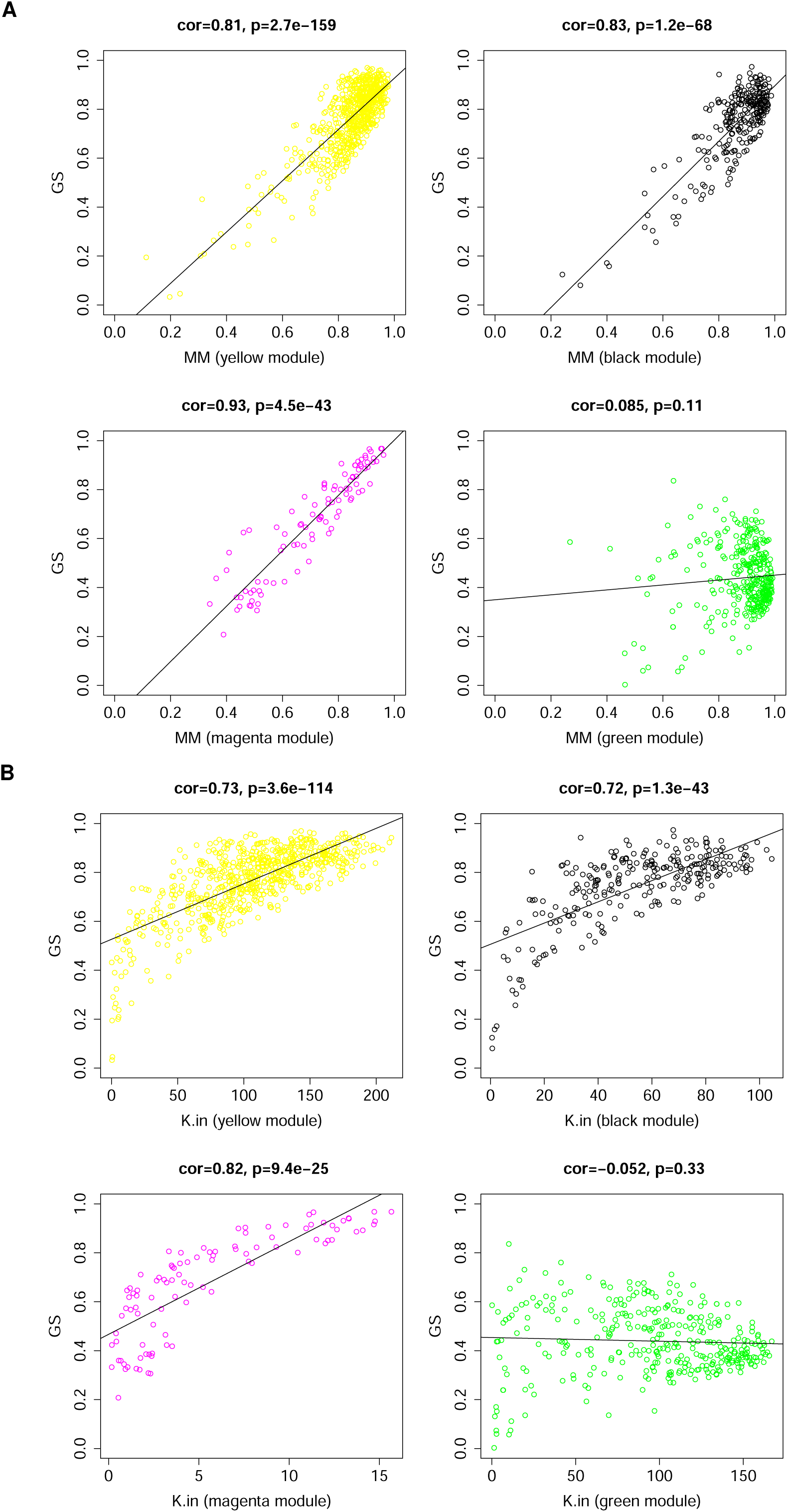
Module features of GS, MM and K.in. (A) Module Membership (MM) and Gene significance (GS) are significantly correlated in the key modules (magenta, black, and yellow). Each point represented an individual gene within each module, which was plotted by GS on the y-axis and MM on the x-axis. The regression line, correlation value, and p-value were shown for each plot. (B) Correlation of the K.in (x-axis) and the GS (y-axis). Green module is not a key module and was added here for sake of comparison.

### Gene hub detection and visualization of module networks

Genes in the selected key modules were ranked according to the intramodular connectivity and the top 20 genes of each key modules was used to visualize the network of each specific module (Fig. 4). Subsequently, the top three genes of the yellow, black, and magenta modules were labeled as the hub genes in their modules that are important for COVID19 disease. Thus, the protein coding genes COPE, CDC37 and CAPNS1 were selected as the hub genes in the yellow module, whereas AURKAIP1, LAMTOR2, and GABARAP protein coding genes were selected as the hub genes in the black module. Regarding the magenta module, MT-ND1, MT-ND5, and MTRNR2L12 are selected as hub genes. All of these hub genes exhibited a high intramodular connectivity, which established their network centrality and potentially vital roles in the COVID19 disease. We also observed that not all of hub genes have shown differential gene expression (Table 1).

**Table 1:** Differential expression of hub genes identified in the key modules. (LogFC: log2 transformed fold change; AveExpr: Average expression; Adj.P.Val: Adjusted P-value).

| Gene | LogFC | AveExpr | Adj.P.Val | Module |
| --- | --- | --- | --- | --- |
| MT-ND5 | -3.2 | 14.0 | $2.12 \times 10^{-5}$ | magenta |
| MT-ND1 | -3.1 | 15.2 | $2.32 \times 10^{-6}$ | magenta |
| MTRNR2L12 | -3.1 | 15.5 | $5.83 \times 10^{-7}$ | magenta |
| AURKAIP1 | 1.6 | 3.1 | $4.29 \times 10^{-3}$ | black |
| GABARAP | 1.2 | 6.4 | $2.75 \times 10^{-3}$ | black |
| COPE | 0.9 | 4.5 | $3.04 \times 10^{-2}$ | yellow |
| LAMTOR2 | 0.9 | 3.4 | $4.08 \times 10^{-2}$ | black |
| CAPNS1 | 0.6 | 8.3 | $1.14 \times 10^{-1}$ | yellow |
| CDC37 | 0.6 | 6.9 | $1.33 \times 10^{-1}$ | yellow |

**Figure 4:**
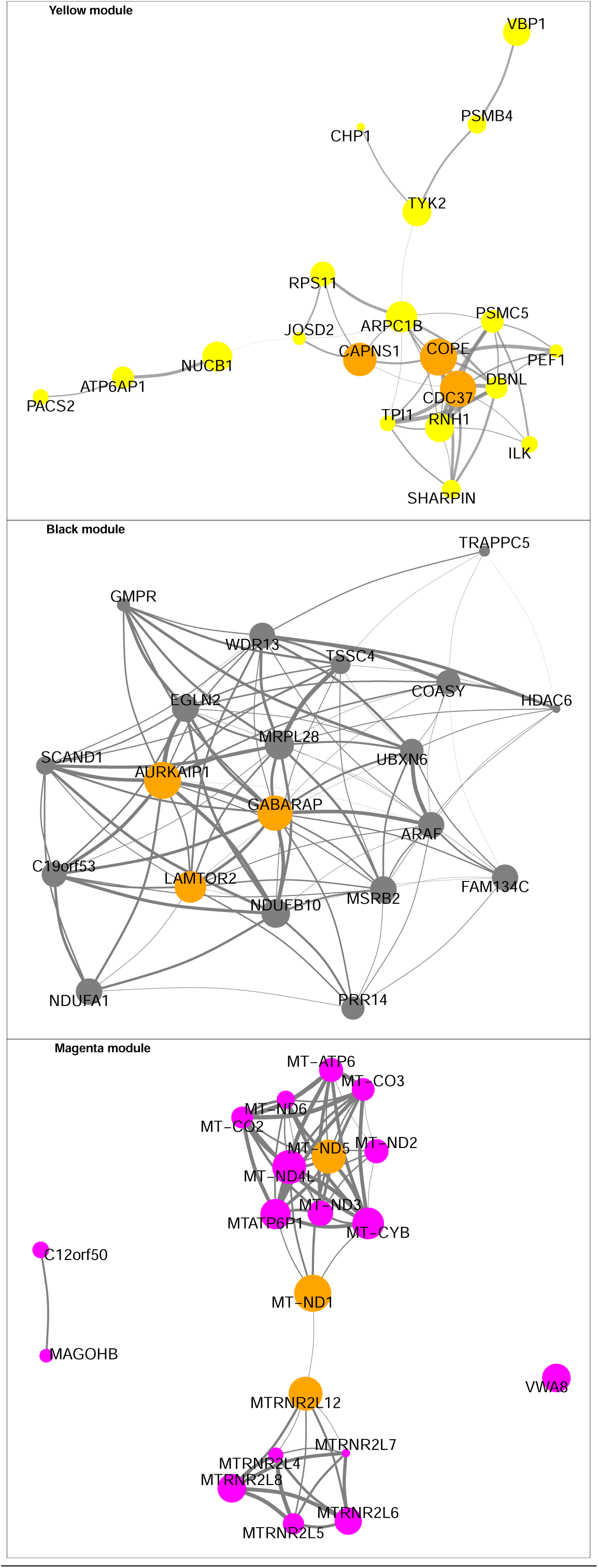
Interaction of gene co-expression patterns in the key identified module and hub gene abundance. The module was visualized using R package “ggraph” software. The node size corresponds to the K.in level. and the thickness of the link represents the strength of correlation between genes. For sake of visualiztion clarity, edges of weight less than 0.6 were not drawn.

### Enrichment Analysis of key modules

Gene ontology (GO) pathway enrichment analyses were performed on the yellow, black, and magenta modules using Targetmine platform, and the top relevant terms of each category are presented in (Fig. 5A). The pathway enrichment results demonstrated that the genes in both yellow and black modules were primarily enriched in pathways associated with metabolic process, protein translation, energy substance metabolism, mitochondrial activity, and oxidative phosphorylation. Genes in the magenta module were enriched in several pathways that are primarily associated with regulation of megakaryocyte differentiation and apoptosis, including the regulation of the execution phase of apoptosis. Reactome showed enriched pathways of metabolism, platelet degranulation, and response to elevated platelet cytosolic Ca^2+^ in the yellow module. The black module shows enrichment of respiratory electron transport, ATP synthesis by chemiosmotic coupling, heat production by uncoupling proteins, citric acid (TCA) cycle, and respiratory electron transport just to name a few (Fig. 5B) (More detailed results are shown in supplementary Fig. 3).

**Figure 5:**
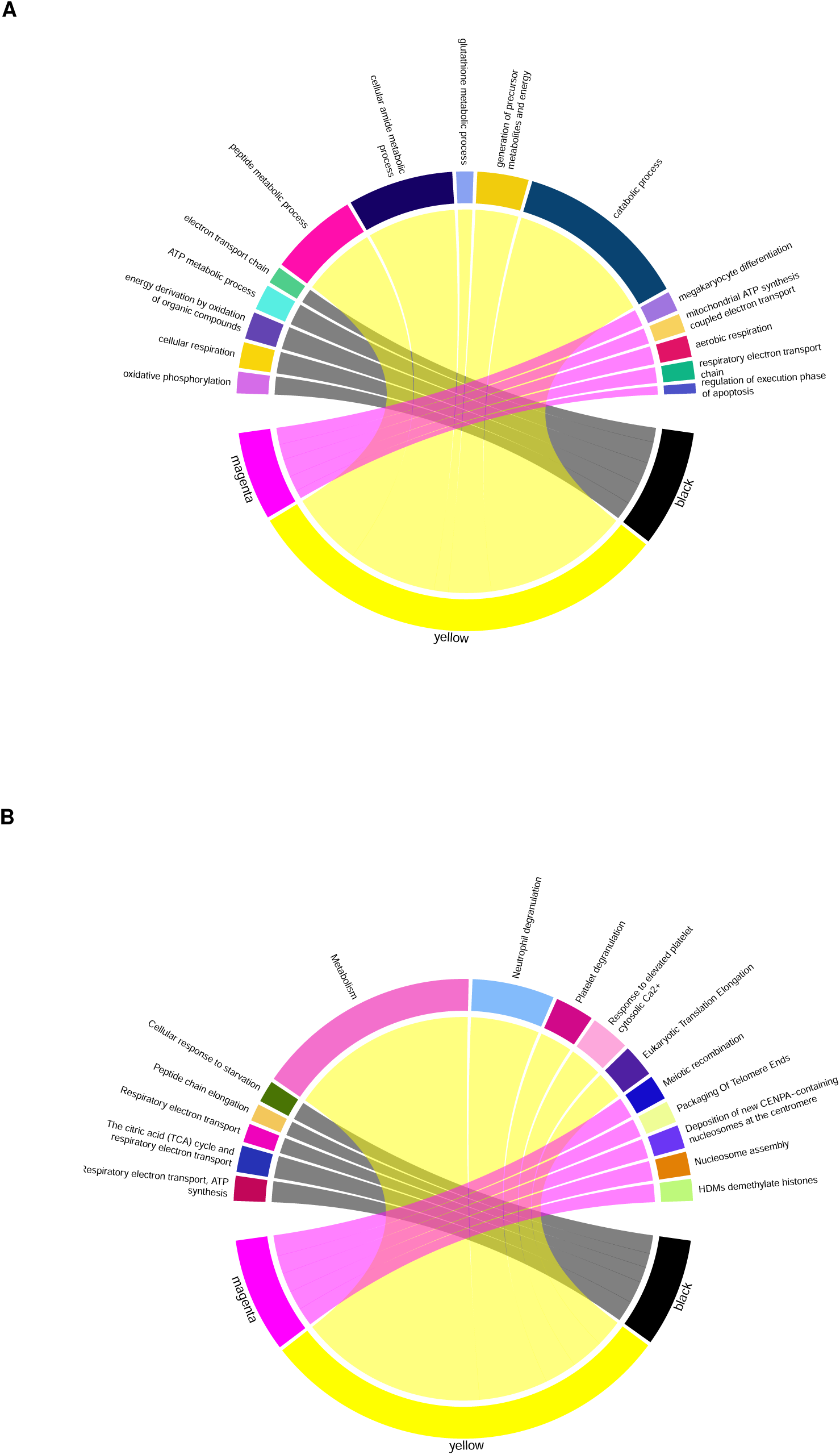
Gene ontology (GO) terms and Reactome enrichment analysis of the key modules. Chord plots depicting the enriched top significant GO terms of biological process (A), and top significant Reactome pathways (B) in the key modules. Thickness of the connection is corresponding to the number of involved genes.

## Discussion

The underlying pathophysiological mechanisms of thrombosis in COVID19 are extremely complicated^32^, and hence clearly require more examination. Inspecting gene co-expression patterns is proven to be an effective method to analyze and uncover complicated genetic networks. To address the aforementioned issues, in the present study, gene co-expression analysis was performed on COVID19 platelet RNAseq dataset containing gene expression data from ten COVID19 patients and five healthy controls. There were three modules that were identified as the key modules in COVID19, with the highest level of significant association. The top three genes of each key module with the highest intramodular connectivity were identified as hub genes for COVID19 in platelets. The results of the enrichment analysis suggest that the key modules and the pathological processes underlying the disease are associated with energy metabolism, mitochondrial process, and apoptosis. Furthermore, we also saw enrichment of platelet secretion and activation pathways. These results provide-at least in partan illustration of the comprehensive platelet regulatory network in COVID19, which should improve the current understanding of the mechanisms underlying immunothrombosis in COVID19 patients. Ultimately, these findings might help in finding appropriate therapeutic targets.

The present study used the data in BioProject accession #PRJNA634489^7^ to perform the co-expression analysis using WGCNA. The data used in this study, which was generated by Manne et al ^7^, revealed that COVID19 disease leads to changes in platelet transcriptional profiles in comparison to control. Manne et al showed that platelet differential gene expression in COVID19 is associated with enrichment of protein ubiquitination, antigen presentation, and mitochondrial dysfunction. The major differences in the genes or modules obtained in the present study, compared with the results from other studies including the one by Manne et. al.^7^ is that the present study used a more comprehensive method by employing WGCNA. Using this method, we were able to identify/pull-out co-expression modules of genes, namely the yellow and black modules, which represent an important regulatory modules of platelet function in COVID19. In addition, we were able to identify the magenta module, which represents genes that are negatively correlated with COVID19 disease state and show enrichment in megakaryocyte differentiation and apoptotic pathways. This systematic and in-depth analysis performed should complement results obtained from conventional DEGs analysis, and therefore allow for a better understanding of the pathophysiological mechanisms of COVID19 disease.

Notably, the co-expression analysis revealed a total cluster of 16 modules, with the yellow and black modules exhibiting the strongest positive correlation with COVID19 disease. Enrichment analysis indicated that the genes in the yellow and black modules were primarily associated with platelet metabolism, energy, and oxidative phosphorylation. Furthermore, the analysis of the yellow module showed enrichment of a host of platelet functional responses/activities, such as platelet degranulation/secretion and increased platelet response to Ca^2+^. Indeed, other studies showed the COVID19 disease to be associated with platelet activation and to increase platelet alpha granule secretion, which are critical in the development of thrombosis seen in those patients^7,8^. It is noteworthy that the platelet alpha granule secretion response is not only important for thrombus formation, but also in inflammation by releasing receptors that facilitate adhesion of platelets with other vascular cells as well as releasing a wide range of inflammatory chemokines^33^.

The yellow and black modules show strong enrichment in platelet metabolic processes, which is in agreement with the increase in platelet activation. For instance, previous data have shown that platelet transition from inactive to active state requires alteration in ATP availability^34^, furthermore, substrate metabolism (e.g. glucose) was shown to be essential for platelet activation^35^, and thrombosis^36^ This seems to suggest that altered platelet metabolism may play a critical role in the pathophysiology of thrombosis in COVID19 patients. It is important to note that reports have suggested that a state of hypermetabolic demand is one of COVID19 disease features, especially when sepsis develops ^37^. Like other viruses that can impact cellular metabolism in human cells and utilize them to their advantage, SARS-CoV-2 virus appears to have the ability to localize proteins to mitochondria and hijack the host’s mitochondrial function ^38^. This mechanism might explain the enrichment of platelet mitochondrial processes we observed in the yellow and black modules. This finding is in fact supported by a recent study that reported that SARS-CoV-2 impacts mitochondria in platelets, which affects their involvement in the pathophysiology of thrombosis in COVID19 patients^39^. The enrichment of protein translation in the yellow and black modules suggests an alteration in protein synthesis and possible hijacking of the translation machinery of platelet by the virus. In line with this observation, one study suggested that the cells infected with SARS-CoV-2 might exhibit a faster protein synthesis rate, which implies a high translation rate ^40^. This notion requires further investigation to determine the exact mechanism underlying enhancement of translation in platelets of COVID19 patient.

One particular characteristic of platelet apoptotic process is the phosphatidylserine (PS) exposure, which is essential for the generation of thrombin^41^. PS exposure is found to be downregulated in activated platelets from COVID19 patients due to mitochondrial dysfunction^39^. This observation is supported by the negative regulation of apoptotic processes in platelets enrichment in the negatively correlated magenta module. On the contrary, another report showed that COVID19 increases PS externalization, which is linked to thrombosis^42^. The impact of platelets mitochondrial damage on hemostasis depends on its severity. Thus, it leads to bleeding by progressing toward apoptosis if it is severe; or toward platelet activation pathways and development of thrombosis risk in case of mild damage^43^. Based on this reasoning, COVID19 disease-caused mitochondrial damage in platelets is probably mild; and hence the thrombotic phenotype still prevails in these patients. Based on these considerations, more investigation is needed to confirm these observations and to understand the underlying mechanisms.

We identified hub genes in each of the key modules. For example, in the yellow module the COPE, CDC37, and CAPNS1, which are protein coding genes involved in vesicle-mediated transport, positive regulation of cellular processes, and regulation of interferons. Furthermore, some of these protein coding genes have also been investigated in platelets and shown to regulate important aspects of their function^44–47^, Interestingly, although our co-expression analysis showed that CAPNS1 is an important hub gene in the yellow module, this gene was not differentially expressed in our differential gene expression analysis. Furthermore, CAPNS1 has shown to play a significant role in regulating platelet activity and thrombosis under hypoxia ^46^, a condition commonly seen in severe COVID19 patients^48^. This observation might indicate that some of the important genes in establishing thrombotic phenotype in COVID19 may not necessarily be differentially expressed.

The hub genes of the black module, AURKAIP1, LAMTOR2, and GABARAP are linked to regulation of mitochondrial activity, regulation of signaling processes, and protein targeting. Data on the role of these genes in platelets is limited, however, further investigation is warranted based on our findings. It is noteworthy that LAMTOR2 is a known regulator of the MAPK/Erk and mTor signaling pathways^49,50^, both of which are shown to be important in regulating platelet function ^51,52^. Moreover, the p14/LAMTOR2 deficiency-which is associated with one of the primary immunodeficiency diseases that also include “Hermansky–Pudlak syndrome type 2”-has been linked to platelet defects^53^. However, more needs to be done to examine the exact role of LAMTOR2 in platelets of COVID19 patients.

In the magenta module, MT-ND1, MT-ND5, and MTRNR2L12 protein coding genes are related to NADH dehydrogenase activity and apoptotic processes. According to our analysis, all hub genes in the magenta module have differentially expressed and are downregulated in COVID19 patients in comparison to healthy controls. Down regulation of MT-ND1, MT-ND5 protein coding gene might at least in part explain the mitochondrial dysfunction seen in platelets of COVID19 patients. With respect to MTRNR2L12, it was observed that it is one of the differentially expressed genes in bronchoalveolar lavage fluid samples from patients with severe COVID19 in comparison to control ^54^. MTRNR2L12 is a paralog of the protein coding gene MTRNR2L8, and both are expressed in platelets ^55^. It is of note that MTRNR2L12 was shown to be among the top 10 RNA with differential splice junctions in platelets of patients of multiple sclerosis ^56^.

In addition to the identified hub genes, a number of other canonical platelet genes in the yellow and black modules were also associated with platelet function. For example, SLEB and ITGA2B protein coding genes were present in the yellow module with high intramodular connectivity (ranked in the top 50) and both proteins are critical for platelet function^57^. Moreover, another canonical platelet gene that was also identified in the yellow module, namely ITGB3 was ranked 132 with regard to its intramodular connectivity, which is considered high in the yellow module of 681 genes. Furthermore, we also noticed that the protein coding gene IFITM3 shows high module membership (black module). The protein encoded by this gene is an interferon-induced membrane protein that was shown to be helpful in immunity against influenza A H1N1 virus, West Nile virus, and dengue virus^58–60^. Most recently, IFITM3 has also been shown to be upregulated protein in COVID19 disease^7,61^.

The present study has certain limitations that should be noted. Firstly, the analysis focused on only one dataset, due to limited access to platelet gene expression data that were collected from COVID19 patients. Therefore, additional datasets should be analyzed, if available, to validate our findings and/or obtain more representative results. Also, the number of samples was 15, which may be associated with some noise, albeit it is the minimum number of samples recommended for co-expression analysis by WGCNA. Finally, any limitations in the original study, from which the data was obtained will also be reflected in the results of this study.

In conclusion, our co-expression analysis of a platelet RNAseq dataset from COVID19 patients and healthy controls revealed 16 modules, amongst which the yellow, black, and magenta were identified as the most critical in COVID19 disease. Additionally, 9 hub genes were determined to potentially serve key roles in pathophysiological mechanisms of COVID19 in the context of platelet biology. The positively associated yellow and black modules were identified to be involved in platelet degranulation, energy metabolism, and mitochondria. The negatively associated magenta module was associated with interactive pathways of apoptosis. These data should help expand our understanding of the underlying mechanisms of thrombosis in COVID19 disease and help promote and guide future experimental studies to investigate the roles of the protein coding genes in the pathophysiology of this disease. Additionally, these genes may serve as novel therapeutic targets for treating patients.

## Acknowledgements

The authors acknowledge Matthew T. Rondina and Robert A. Campbell from the department of internal medicine, University of Utah, Salt Lake City, Utah, USA, for their contribution and suggestions that improved the work substantially. The statements made here are solely the responsibility of the authors. No external funding was utilized in this study.

## Authorship

### Contribution

AA, AM: conceptualization; AA, AM: analyzed results and made the figures; AA: drafted the manuscript; AA, AM, KM, FA, FK: edited the manuscript and approved the submission;

## Supplementary Material

**Supplementary Figure 1:**
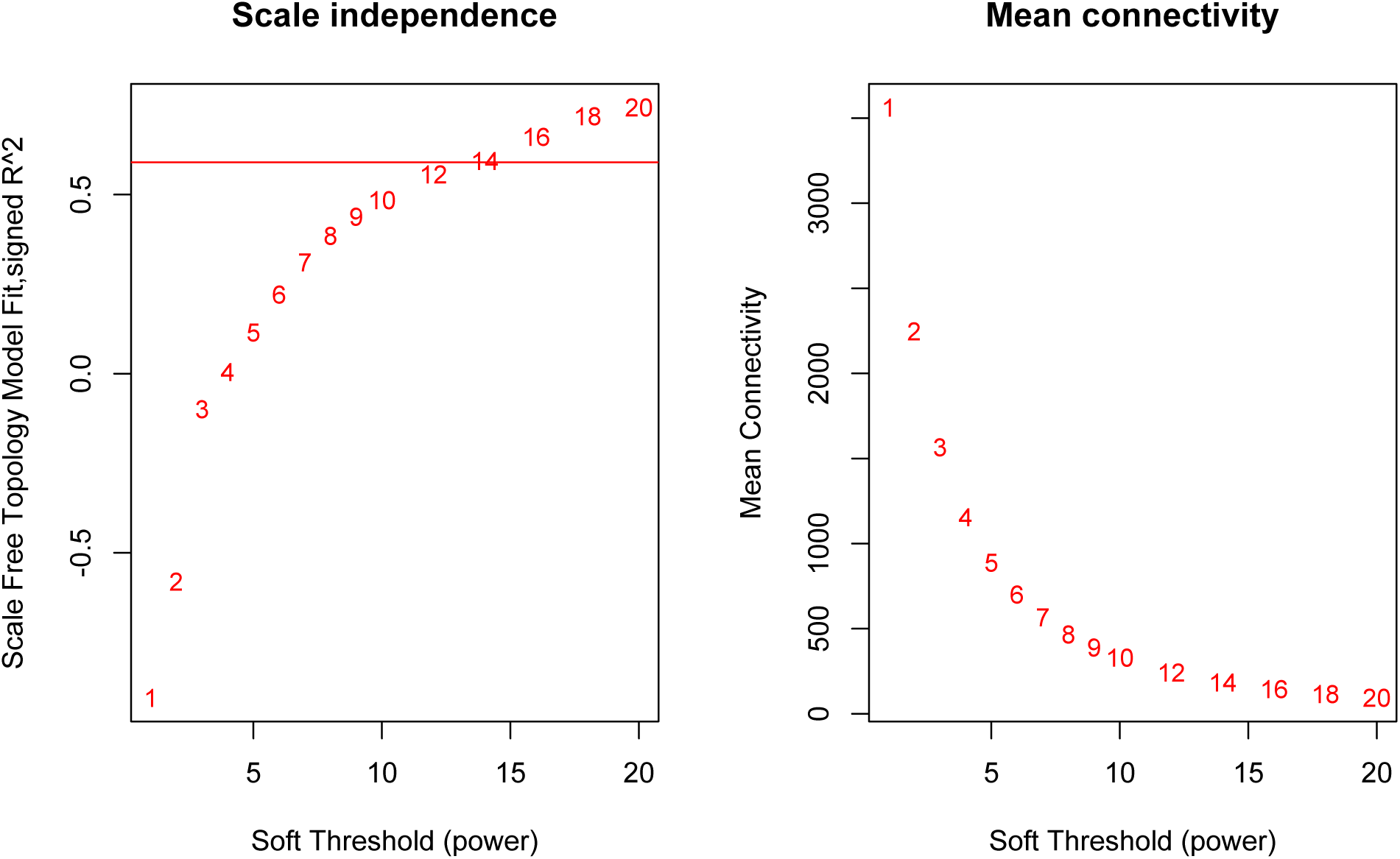
Network Soft threshold

**Supplementary Figure 2:**
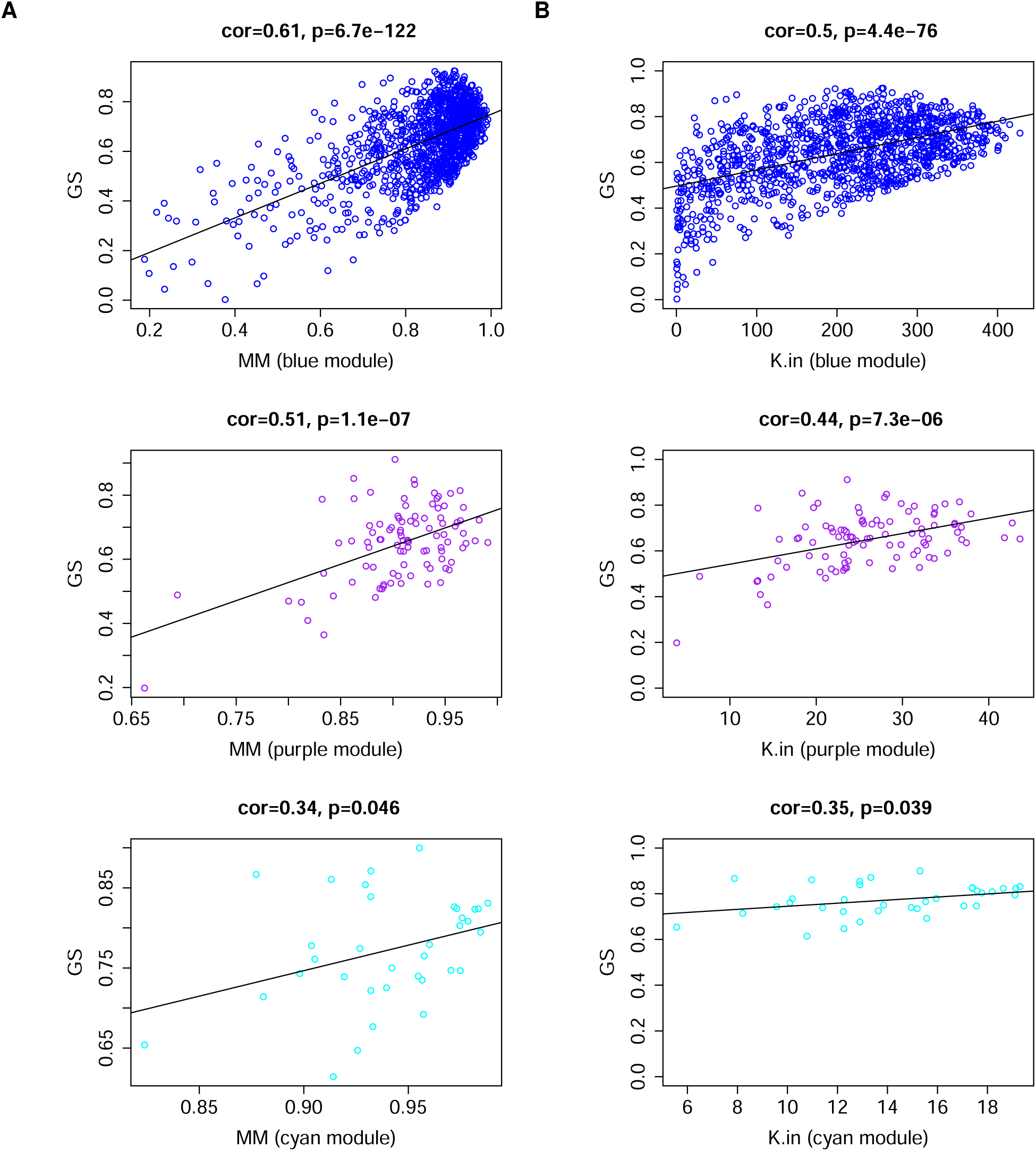
Module Membership and gene significance in other modules.

**Supplementary Figure 3:**
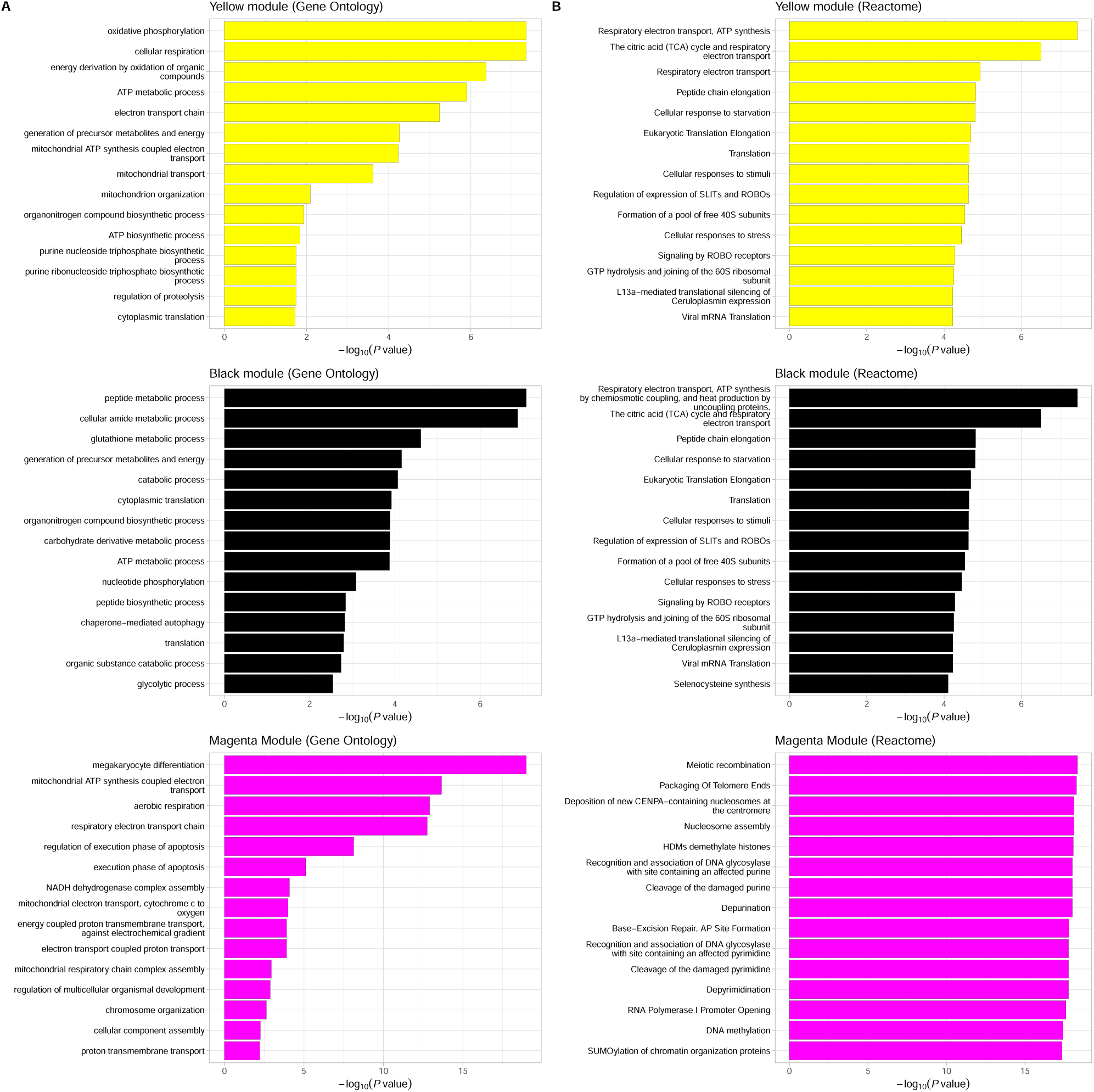
Detailed enrichment analysis. A; Gene Ontology of biological function. B; Reactome pathways enrichment.

